# The K-Cl co-transporter 2 is a point of convergence for multiple autism spectrum disorder and epilepsy risk gene products

**DOI:** 10.1101/2020.03.02.973859

**Authors:** Joshua L. Smalley, Georgina Kontou, Catherine Choi, Qiu Ren, David Albrecht, Krithika Abiraman, Miguel A. Rodriguez Santos, Christopher E. Bope, Tarek Z. Deeb, Paul A. Davies, Nicholas J. Brandon, Stephen J. Moss

## Abstract

KCC2 plays a critical role in determining the efficacy of synaptic inhibition and deficits in its activity lead to epilepsy and neurodevelopmental delay. Here we use unbiased proteomic analyses to demonstrate that KCC2 forms stable protein complexes in the neuronal plasma membrane with 96 autism and/or epilepsy risk gene (ASD/Epi) products including ANKB, ANKG, CNTN1, ITPR1, NCKAP1, SCN2A, SHANK3, SPTAN1, and SPTBN1. Many of these proteins are also targets of Fragile-X mental retardation protein (FMRP), the inactivation of which is the leading monogenic cause of autism. Accordingly, the expression of a subset of these KCC2-binding partners was decreased in Fmr1 knockout mice. Fmr1 knockout compromised KCC2 phosphorylation, a key regulatory mechanism for transporter activity and the postnatal development of GABAergic inhibition. Thus, KCC2 is a point of convergence for multiple ASD/Epi risk genes and therapies targeting this transporter may have broad utility in alleviating these heterogeneous disorders and their associated epilepsies.

---

The K_+_/Cl_−_ co-transporter KCC2 (encoded by the gene SLC12A5) is the principal Cl_−_-extrusion mechanism employed by mature neurons in the CNS (1). Its activity is a pre-requisite for the efficacy of fast synaptic inhibition mediated by Glycine (GLYR) and type A γ-aminobutyric acid receptors (GABA_A_R), which are Cl_−_ permeable ligand-gated ion channels. At prenatal and early postnatal stages in rodents, neurons have elevated intracellular Cl_−_ levels resulting in depolarizing GABA_A_-mediated currents (2). The postnatal development of canonical hyperpolarizing GABA_A_R currents is a reflection of the progressive decrease of intraneuronal Cl_−_ levels that is caused by the upregulation of KCC2 expression and subsequent activity (3–6). These changes in neuronal Cl_−_ extrusion reflect a sustained increase in the expression levels of KCC2 after birth, the mRNA levels of which do not reach their maximal levels in humans until 20-25 years of age (7). In addition to this, the appropriate developmental appearance of hyperpolarizing GABA_A_R currents is also in part determined by the phosphorylation status of KCC2, a process that facilitates its membrane trafficking and surface activity (8–13).

In keeping with its essential role in determining the efficacy of synaptic inhibition, humans with mutations in KCC2 develop severe epilepsy soon after birth (14–16). Deficits in KCC2 activity are also believed to contribute to the development of temporal lobe epilepsy (17, 18), in addition to other traumas including ischemia and neuropathic pain (19, 20). Given the critical role that KCC2 plays in determining the maturation of inhibitory neurotransmission, subtle changes in its function are also strongly implicated in autism spectrum disorders (ASD) (21). The most common monogenic cause of ASDs is the inactivation of the fragile X-mental retardation protein (FMRP). Inactivation of FMRP in mice (FMR1 KO), a widely used model of autism, delays the postnatal development of GABAergic inhibition. Likewise, alterations in KCC2 function are associated with Downs syndrome (22), fragile X syndrome (23), and Rett syndrome (24–26).

Accumulating evidence from high-throughput sequencing of patients has identified a divergent list of >400 ASD and epilepsy risk genes (ASD/Epi), many of which have been replicated in murine models (27). These genes encode proteins with a broad array of functions, however, the mechanisms by which these heterogenous gene products exert their pathological effects remains ill-defined (28). Here, we isolated stable protein complexes that contain KCC2 from the plasma membrane of mouse forebrain, and characterized their composition using an unbiased proteomic approach. In this way we identified many novel KCC2-associated proteins. Using bioinformatic analyses we discovered that KCC2 forms stable protein complexes containing many ASD/Epi risk gene products, several of which are classed as ‘high-risk’ (29). We then demonstrated that a selected cohort of high-risk proteins, including KCC2 itself, are aberrantly expressed in plasma membrane fractions from FMR1 KO mice, correlating with altered KCC2 phosphorylation at known and novel sites. Thus, KCC2 is a point of convergence for multiple ASD/Epi risk genes that act in part to regulate its phosphorylation.

## Results

### Isolating native stable protein complexes containing KCC2 from brain plasma membranes

In order to define which proteins are associated with KCC2 in neuronal plasma membrane we developed a novel methodology for isolating stable multiprotein complexes enriched in this transporter. To do so, fresh isolated mouse forebrain was homogenized in detergent-free conditions to preserve organelle structure. The resulting homogenate was then subjected to differential gradient-based centrifugation and the plasma membrane fraction isolated (Figure 1a). To confirm effective fractionation, we immunoblotted for established protein markers for each specific organelle (Figure 1b); HSP90 (cytosol), HSP60 (mitochondria), calreticulin (ER), and n-cadherin (n-Cadh) (plasma membrane). Each organelle fraction showed enrichment for its respective marker protein. Importantly, the plasma membrane fraction was both enriched for n-Cadh and depleted for HSP60, demonstrating efficient separation of plasma membrane and mitochondria, a common contaminant of membranous fractions. KCC2 was enriched in both the mitochondrial and plasma membrane fraction. The enrichment of KCC2 in our target fraction - the plasma membrane fraction, was highly encouraging for downstream purification. The presence of KCC2 in the mitochondrial fraction could be an interesting future area of research, as chloride homeostasis fundamental for correct mitochondrial function.

**Figure 1.**
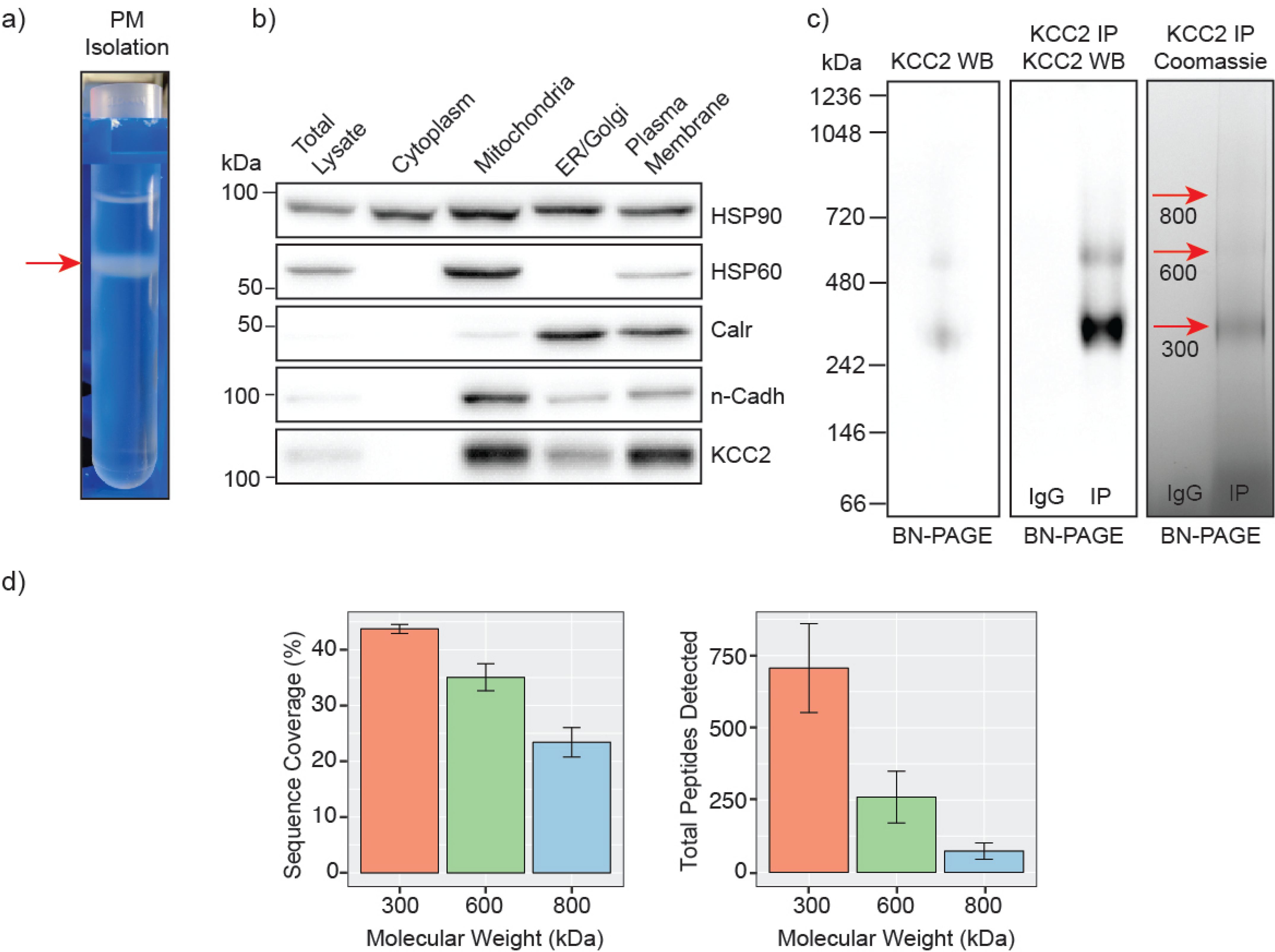
Highly enriched KCC2-containing protein complexes were isolated from purified plasma membranes from mouse forebrain. **A.** Differential centrifugation of homogenized forebrain from 8-12-week-old mice was used to fractionate cellular organelle and obtain an enriched plasma membrane fraction. **B.** Western blots of the organelle fractions were used to measure the abundance of organelle markers; HSP90 (cytosol), HSP60 (mitochondria), calreticulin (ER/Golgi), n-Cadherin (n-Cadh) (plasma membrane). KCC2 abundance was also measured to confirm KCC2 enrichment in the plasma membrane fraction. **C.** Blue Native PAGE (BN-PAGE) was carried out on plasma membrane lysates, and immunoprecipitated KCC2 to resolve the native protein complexes that contain KCC2. These were correlated with protein bands observed by Coomassie staining. **D.** Detected peptides were mapped to the KCC2 reference sequence to determine KCC2 sequence coverage obtained from LC-MS/MS of BN-PAGE protein bands produced following KCC2 IP. Bar charts of the KCC2 sequence coverage expressed as a percentage of the full sequence and by total KCC2 peptides detected (n=4).

Following detergent solubilization, membrane fractions were subject to immunoprecipitation with a monoclonal antibody directed against the intracellular C-terminus of KCC2 covalently coupled to protein G-coated ferric beads. Initial optimization experiments revealed that we were able to isolate approximately 65% of the KCC2 available in the lysate (Figure S1a, b), and the lysate preparation produced minimal KCC2 oligomerization when resolved by SDS-PAGE. Purified material was eluted under native conditions using a buffer containing 2%Tween a ‘soft elution buffer’ (30), and subjected to Blue-Native-PAGE (BN-PAGE; Figure 1c). We observed well-resolved bands by Coomassie staining at 300, 600, and 800 kDa. These bands were not seen in control experiments performed with non-immune mouse IgG. An immunoblot of the same sample confirmed that these bands contained KCC2. Immunoblots of plasma membrane lysates separated by BN-PAGE showed that these distinct KCC2 complex bands exist in the lysate and are not an artifact of or degraded by purification. Despite the high stringency of our method (with high speed centrifugation and low concentration SDS exposure), no monomeric KCC2 was observed (~150 kDa), indicating that KCC2 was maintained in higher order complexes. This may disrupt low affinity protein interactions and favor the detection of higher affinity binding partners.

After we confirmed that these well resolved bands contained KCC2, we assessed their KCC2 content by LC-MS/MS following trypsin digestion (Figure 1d). The 300, 600, and 800 kDa bands contained an average of 706, 260, and 74 total peptides for KCC2 respectively, which equated to coverage of 44%, 35%, and 23% respectively. In all three bands, the majority of peptides corresponded to the cytoplasmic and extracellular domains of KCC2, while peptides corresponding to the transmembrane domains were rarely detected. The failure to detect these regions is consistent with their high hydrophobicity and subsequent low recovery by LC-MS/MS (Figure S2a). We also noted that the 300, 600 and 800 kDa bands contained peptides derived from KCC1, which were at approximately 6-fold lower levels than those derived from KCC2. Taken together, these results demonstrate our experimental methodology facilitates the isolation of stable high molecular mass protein complexes enriched in KCC2.

### Analyzing the composition of stable protein complexes of KCC2

Having confirmed the veracity of our KCC2 purifications we investigated which proteins form stable protein complexes with this transporter in the brain. First, we compiled a data matrix of all the proteins and total peptide counts detected by proteomic analysis in each of our samples (Table S1). We compared the average total peptide counts for KCC2, with the average total peptide counts for all proteins that were detected in our LC-MS/MS spectra (Figure 2a). In the 300, 600, and 800 kDa bands, KCC2 was the most abundant protein detected, however the proportion of peptides for KCC2 relative to peptides for all other detected proteins decreased with increasing complex molecular mass, this is likely due to the increased protein complexity of the high molecular weight complexes.

**Figure 2.**
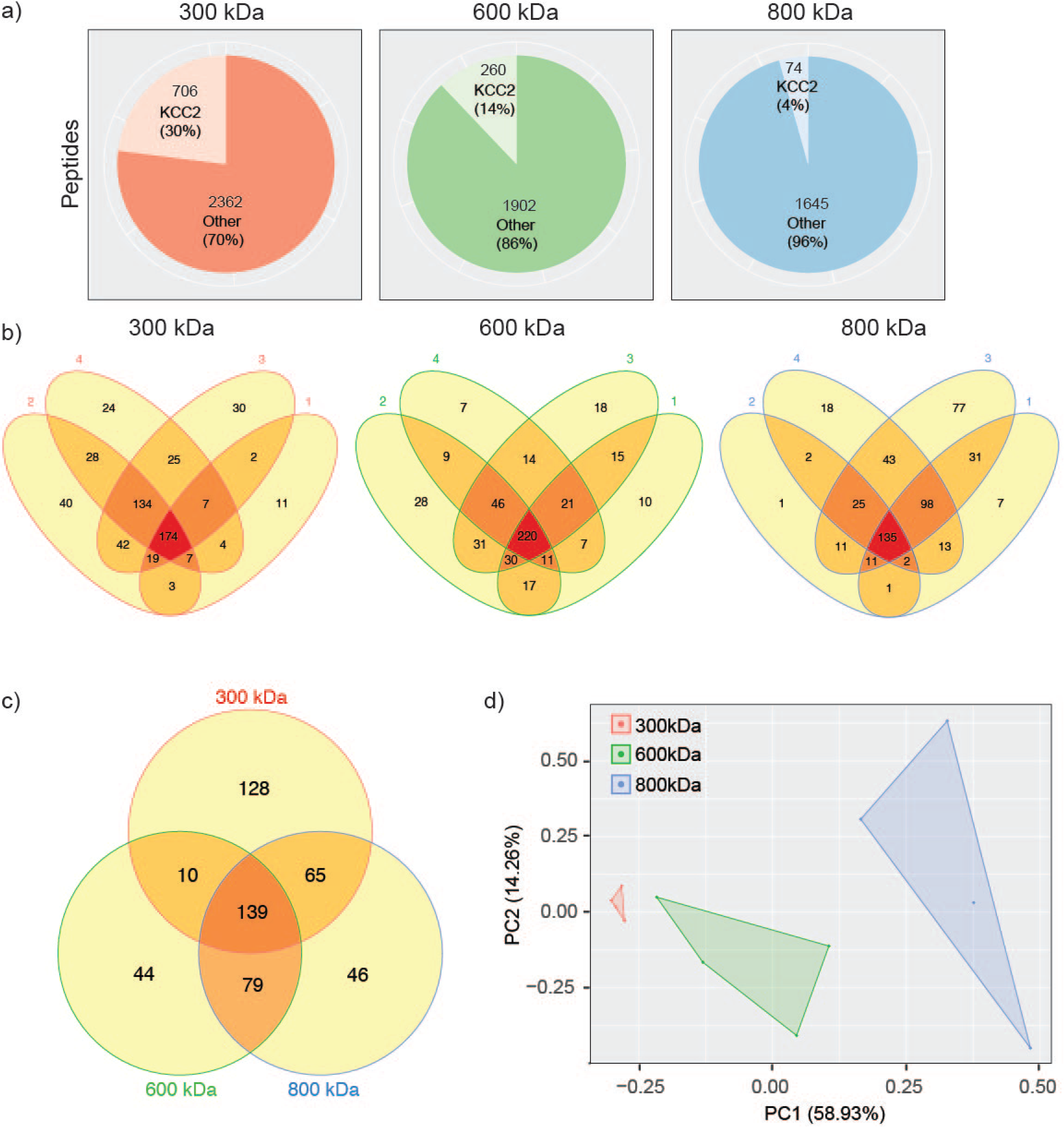
Stable protein complexes contain large amounts of KCC2 and a robust set of associated proteins, which differ subtly between the different molecular weight bands. **A.** Pie charts showing the average number of total KCC2 peptides detected by LC-MS/MS relative to the average number of total peptides detected for all other proteins in each molecular weight complex. **B.** Venn diagrams showing the number and overlap of KCC2-associated proteins identified in each of 4 biological replicates in each molecular weight complex. **C.** Venn diagram showing the overlap in associated proteins detected in at least 3 of the 4 replicates for each molecular weight complex. **D.** PCA analysis of each biological replicate for each molecular weight complex based on their protein composition.

Next, we evaluated the reproducibility of the detected proteins in each of the 4 repeats for each band size (Figure 2b). The detected proteins for each band size were highly reproducible with only proteins detected in at least 3 repeats taken forward for further analysis as ‘robust binding proteins.’ Proteins that also were found in control purifications on control IgG were removed as components of the CRAPome common contaminants found in affinity-MS data (31, 32). We then compared these robust binding proteins between the 300, 600, and 800 kDa complexes (Figure 2c). There was a high degree of overlap between the 600 and 800 kDa complexes. The 300 kDa complex had considerable overlap with the 600 and 800 kDa complexes, but it also contained many unique proteins. This unique protein pool is likely to be large proteins that have dissociated from KCC2 complexes.

Finally, we used principle component analysis (PCA) to assess the degree of similarity in binding protein patterns between all of the proteomic samples (Figure 2d). PCA identifies variance between samples and expresses the degree of similarity by proximity in a PCA plot. We did this to both assess the reproducibility of the biological replicates and the degree of difference in the binding proteins for each complex. The binding proteins were assembled into a matrix of all repeats and detected proteins along with total peptide counts for each. The data was normalized by z-transformation (Table S2) and the PCA plotted created using the default settings in the ggfortify package (accessed January 2019) in R (33). The datasets clearly separated into biological repeats of the different mass bands. This demonstrates that we have identified a reproducible set of binding proteins for the 300, 600, and 800 kDa bands. While there is a substantial degree of overlap between the interacting proteins from each molecular mass band, their protein content is different enough that they can be clearly differentiated by PCA. Due to the high number of KCC2 peptides, the degree of unique proteins, and the potential for unbound proteins being present at 300 kDa, we only investigated the 600 and 800 kDa bands further for KCC2 associated proteins.

We subjected the top 150 proteins associated with KCC2 in the 600 and 800 kDa bands to network analysis in order to investigate the known protein interactions within each cohort. This was carried out using data from StringDB for previously described experimental interactions (34). We also carried out Gene Ontology (GO) analysis to determine the most overrepresented biological process terms (34). In this way we were able to visualize KCC2 networks and subnetworks along with developing insights into the functional groups of proteins that associate with KCC2 (Figure 3a and b). Both the 600 and 800 kDa protein groups contained subnetworks of proteins associated with endocytosis (AP2A1, SNAP91, SNAP25, and AAK1), cellular localization (SPTAN1, SPTBN1, ANK2, ANK3, and MYO5A) and ion transport (ATP1A1, ATP1A2, ATP2A2, and ITPR1). Critically, association with the Na_+_/K_+_ ATPase alpha subunits supports previous findings in human expression patterns of the concurrent developmental upregulation of both transporters (7). The 800 kDa complex also contained a prominent subnetwork of proteins associated with synaptic signaling and organization (GABBR1, GABBR2, GRM2, and GRM3). Importantly, the GABAB receptor has previously been demonstrated to interact with KCC2 directly (35), which further substantiates our findings. For approximately half of the proteins in both the 600 and 800 kDa complexes, there were no known interactors other than KCC2.

**Figure 3.**
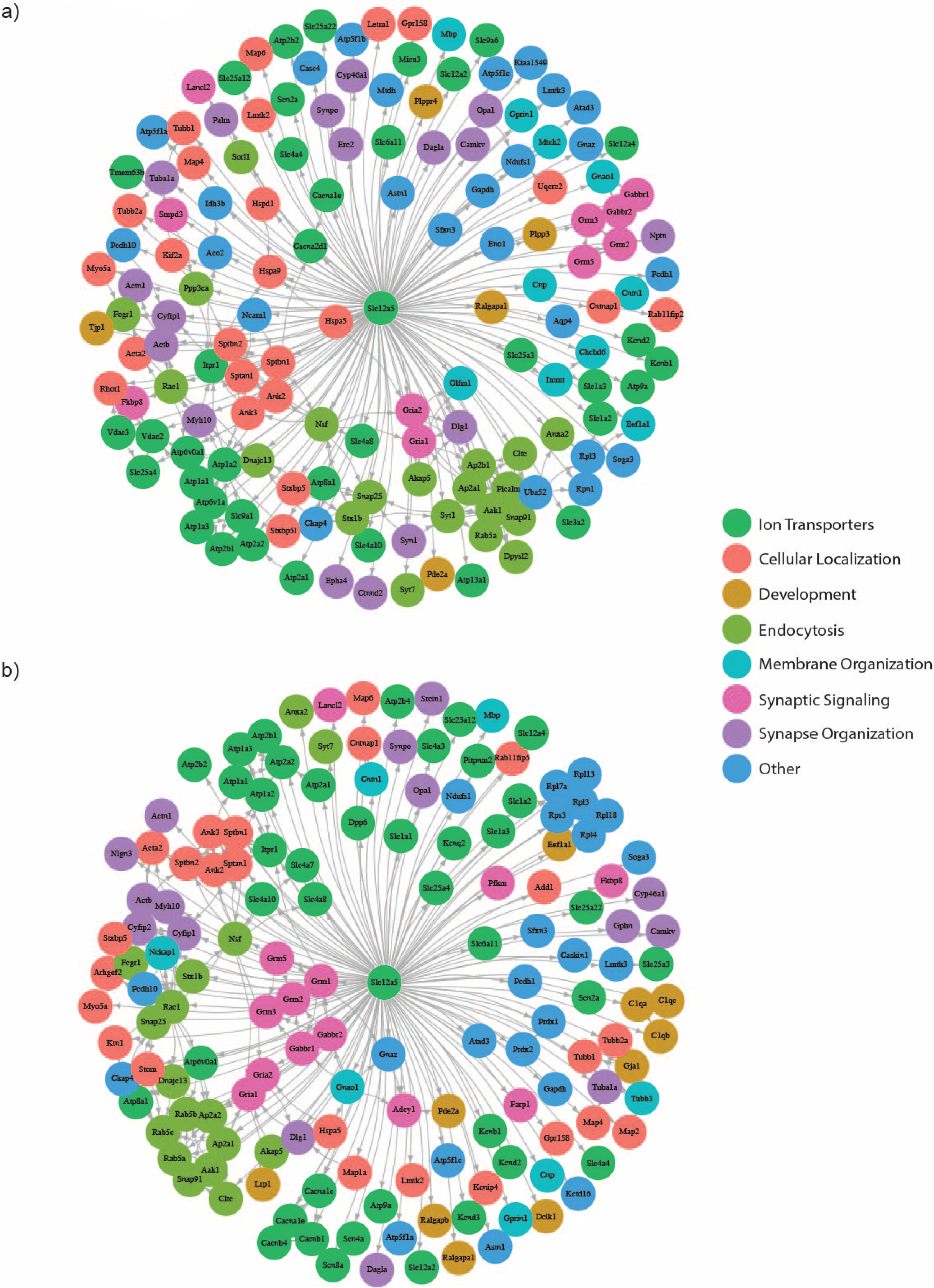
KCC2 is associated with a robust set of proteins from multiple functional classes, with some highly interconnected subnetworks. **A.** A network diagram including the 150 most abundant proteins from the 600 kDa complex based on total peptide counts. Known interactions were obtained using stringent high confidence, direct experimental association parameters from StringDB. These were used to construct a network diagram of protein nodes and arrows to indicate known interactions. The interactions for each protein with KCC2 were included as discovered here. An overlay of gene ontology (GO) terms was used to provide protein classification information. **B.** A network diagram for the most abundant 150 proteins in the 800 kDa complex, analyzed as described above (n=4).

### KCC2 is associated with multiple ASD/Epi risk genes, that are targets for FMR1

As altered KCC2 function has been implicated in ASD and epilepsy, we compiled a list of ASD- and epilepsy-associated genes from SFARI (FMRP targets) (36), and EpilepsyGene (37) (Table S5). We compared this list with the top 150 KCC2-associated proteins in both the 600 and 800 kDa bands. We noted that a number of proteins implicated in either ASD or epilepsy or both co-purified with KCC2 from brain plasma membranes (Figure 4a and b). These comprised several of the highest risk genes associated with autism identified by SFARI, including ANK2, SCN2A (also a major cause of Dravet syndrome) (38), and SHANK3. They also included several products of genes with lower SFARI risk scores, including NCAKP1, MYO5A, ANK3, CACNA1E, CYFIP1, ITPR1 and GPHN, several of which are also highly implicated in various forms of epilepsy. In the 600 kDa complex, 77 out of 150 proteins were ASD/Epi gene products. In the 800 kDa complex this increased to 90 out of 150. In both the 600 and 800 kDa complexes, there are small subnetworks of known interactions between the associated proteins involved in endocytosis and cellular location but for the majority of proteins, KCC2 is the only common factor.

**Figure 4.**
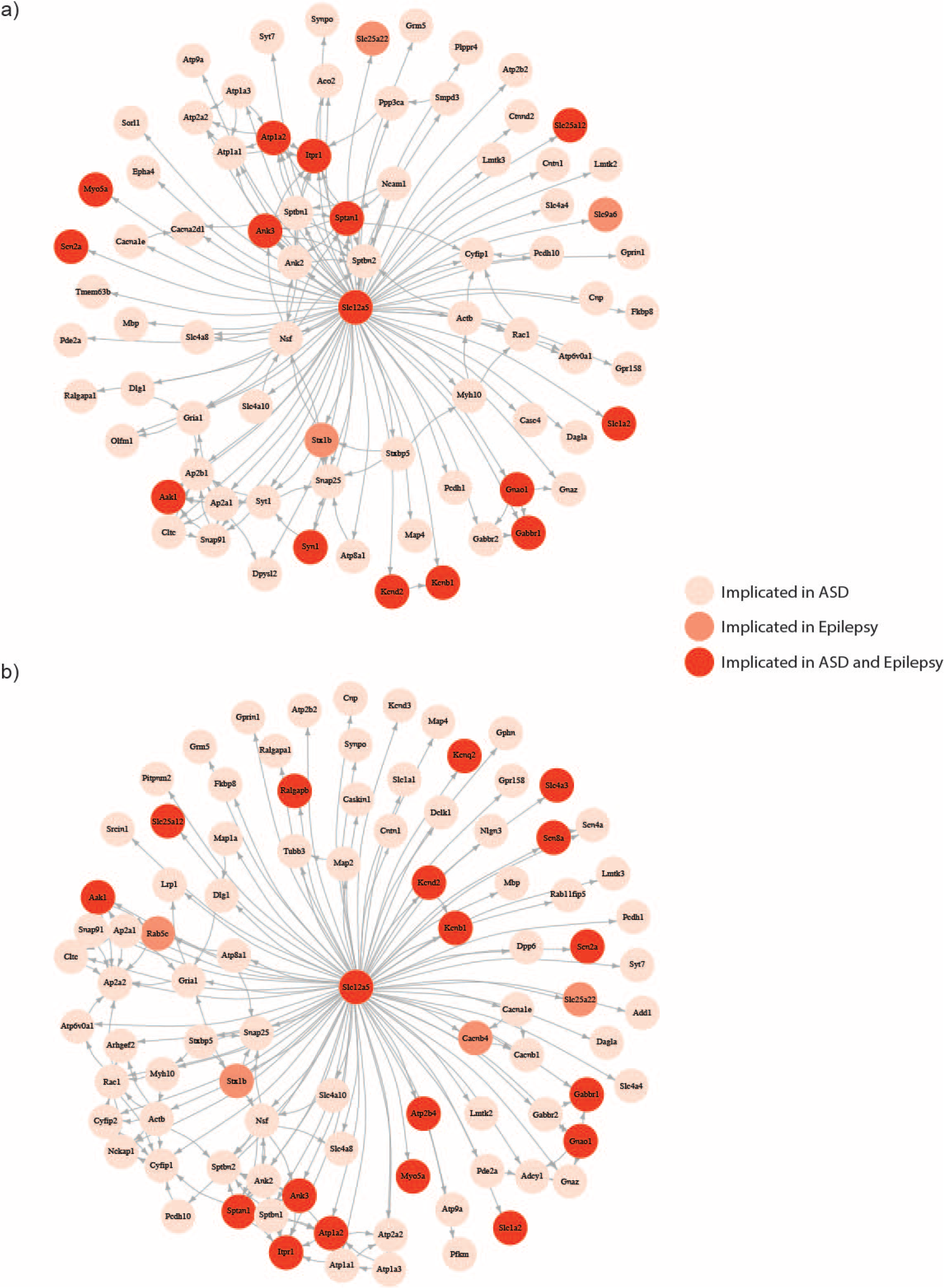
The KCC2 proteome is highly enriched for autism spectrum disorder and epilepsy risk gene products. **A.** A network diagram of KCC2-associated proteins from the 600 kDa complex, that are also the protein products of genes associated with ASD or epilepsy. The known protein associations were sourced from StringDB and applied with an overlay of ASD/Epi risk genes from the SFARI, and EpilepsyGene databases. The colors indicate whether the proteins are implicated in ASD or epilepsy alone, or both. **B.** The same analysis was carried out for the 800 kDa protein complex (n=4).

Next, we set out to confirm that a cohort of high-risk ASD/Epi gene products co-immunoprecipitated in high molecular weight complexes with KCC2 (Figure 5). These proteins were selected by a combination of total peptide hits detected, SFARI risk score, and implication in epilepsy. We immunoblotted isolated KCC2 protein complexes resolved by BN-PAGE for ANK2, ANK3, CNTN1, ITPR1, NCKAP1, SCN2A, SHANK3, SPTAN1, and SPTBN1. No immunoreactivity was observed in the control lanes, whereas the high molecular weight protein complexes of KCC2 showed varying immunoreactivity for all 8 proteins. ANK2, ANK3 (antibody immunoreactive with all splice variants) and NCKAP1 showed specificity for the 600 kDa complex, whereas ITPR1, SCN2A and SPTBN1 showed specificity for the 800 kDa complex. SPTAN1 was equally distributed in the 600 and 800 kDa bands. Only CNTN1 showed positive immunoreactivity for the 300 kDa band. This further demonstrates the specificity of the interactors for high molecular weight KCC2-containing protein complexes.

**Figure 5.**
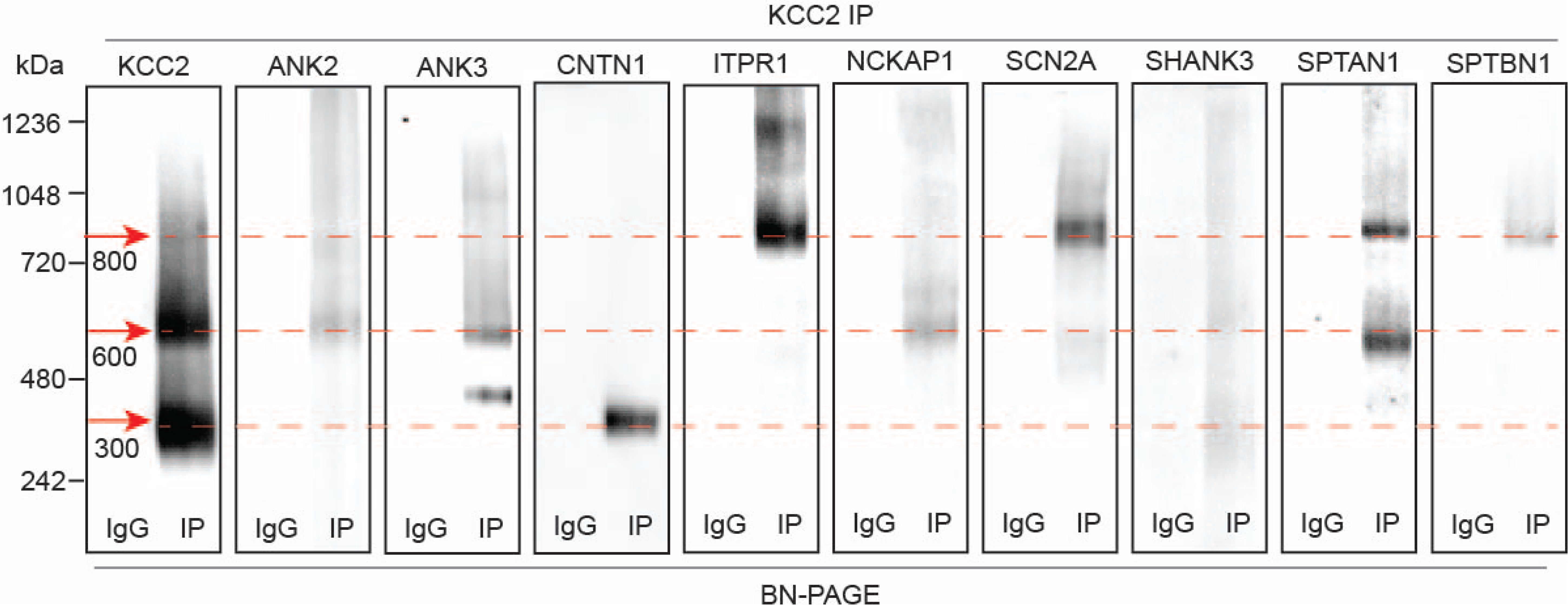
Immunoblots confirm the presence of several high risk ASD/Epi risk gene products in protein complexes that contain KCC2. **A.** KCC2 protein complexes were isolated from forebrain plasma membrane fractions of 8-12-week-old mice, resolved by BN-PAGE and immunoblotted for selected high risk ASD/Epi risk gene products; ANK2, ANK3, CNTN1, ITPR1, NCKAP1, SCN2A, SHANK3, SPTAN1 and SPTBN1.

### KCC2 co-localizes with multiple ASD/Epi proteins on, or close to the neuronal plasma membrane

Having established that KCC2 forms protein complexes with several ASD/Epi risk gene products, we used immunocytochemistry to investigate the proximity of each ASD/Epi risk gene product to KCC2. We used DIV21 mouse primary cultured cortical/hippocampal neurons infected with CamkII-AAV-GFP to ensure we imaged excitatory neurons and to visualize the morphology of the cell and identify whether the interaction between KCC2 and ASD/Epi risk gene products is restricted to specific cellular compartments (Figure 6a and b). SCN2A, ANK3 and NCKAP1 showed punctate staining along dendrites with extensive colocalization with KCC2. This was also evident in the fluorescent density plots by the presence of overlapping peaks. ANKB, CNTN1, ITPR1, and SHANK3 exhibited punctate dendritic staining patterns immediately adjacent to KCC2 puncta. SPTAN1, and SPTBN1 were present in large dendritic clusters with KCC2 mostly located on the edges. Taken together, these data demonstrate that proteins associated in high molecular weight complexes with KCC2 are also highly colocalized with KCC2 in neurons.

**Figure 6.**
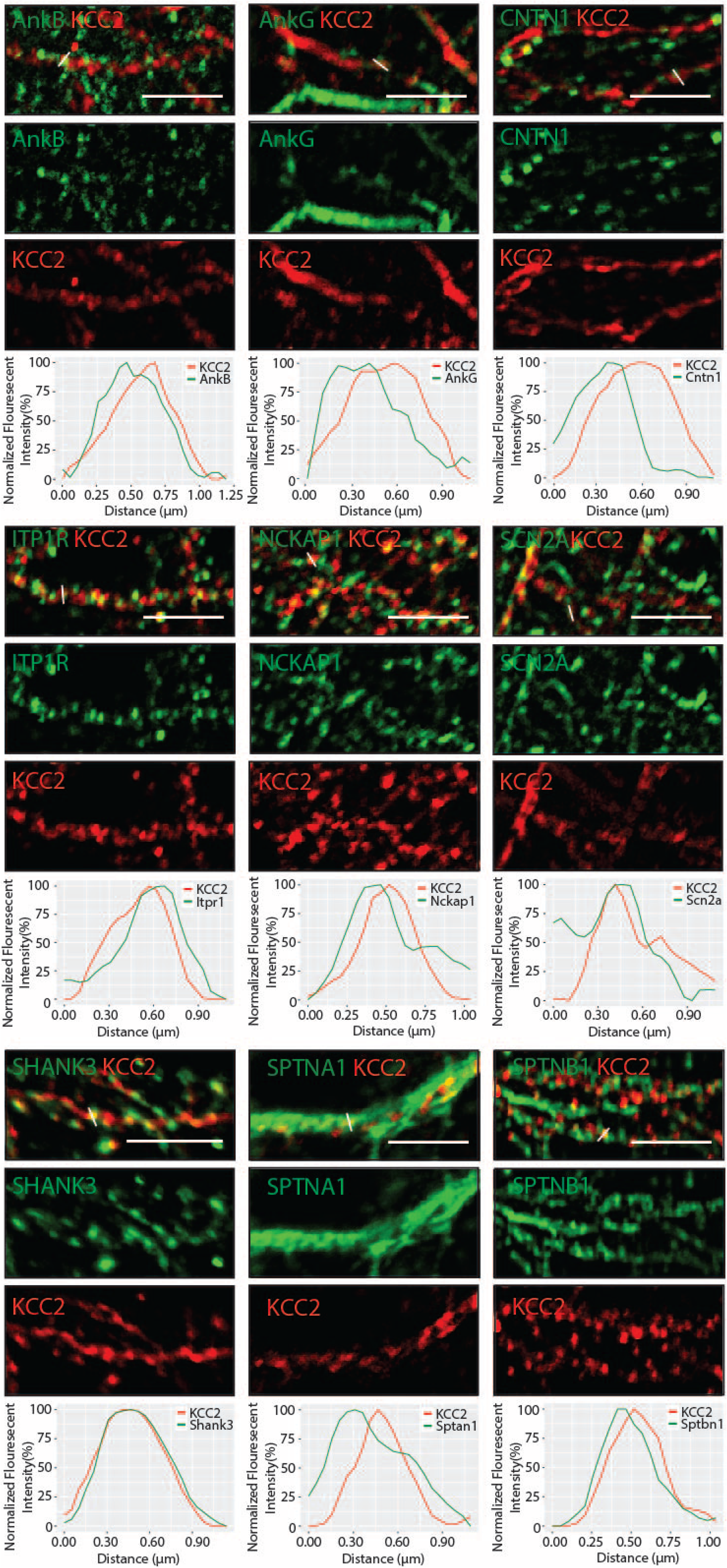
High risk ASD/Epi risk gene products that are in complex with KCC2, colocalize with KCC2 in several neuronal cellular compartments. **A.** Primary cultured neurons from P1 pups were infected with CAMKII AAV-GFP at DIV 3 (to visualize cell morphology and to identify excitatory neurons) and fixed at DIV 21. The cells were immunostained for KCC2 and high risk ASD/Epi risk gene products; ANK2, ANK3, CNTN1, ITPR1, NCKAP1, SCN2A, SHANK3, SPTAN1 and SPTBN1 (n=3).

### Inactivation of Fmr1 reduces the expression of KCC2 and subset of ASD/Epi gene products

We noted that many of the proteins associated with KCC2 are targets for mental retardation protein (FMRP), the mutation of which is the leading monogenic cause of autism. FMRP is an RNA binding protein which regulates the translation of target mRNAs. Thus, we assessed the effects of FMRP on the expression levels of KCC2 and selected components of its proteome using forebrain extracts of Fmr1 knock out mice (FX), a widely used animal model **of** autism. We used immunoblotting to measure the expression levels of proteins of interest in both the total and in the plasma membrane fractions from both genotypes (Figure 7a and b). KCC2 expression levels in the total lysates (TL) were unchanged between wild type (WT) and FX samples. Interestingly there was a significant decrease in KCC2 expression in the plasma membrane fraction (p=0.04). The 190 kDa form of ANK3 was also reduced in the plasma membrane fraction of FX animals (p=0.001), although the 270 kDa and 480 kDa forms were unchanged. CNTN1 expression was reduced in FX mice compared to WT in both the TL (p=0.01) and PM (p=0.01) fractions. Finally, ITPR1 expression in the plasma membrane fractions was also reduced in FX animals (p=0.02). No significant difference in expression was observed in ANK2, NCKAP1, SHANK3, SPTAN1, or SPTBN1. Taken together, these data demonstrate that the expression of several KCC2-associated proteins is reduced in plasma membrane fractions from FX mouse forebrain.

**Figure 7.**
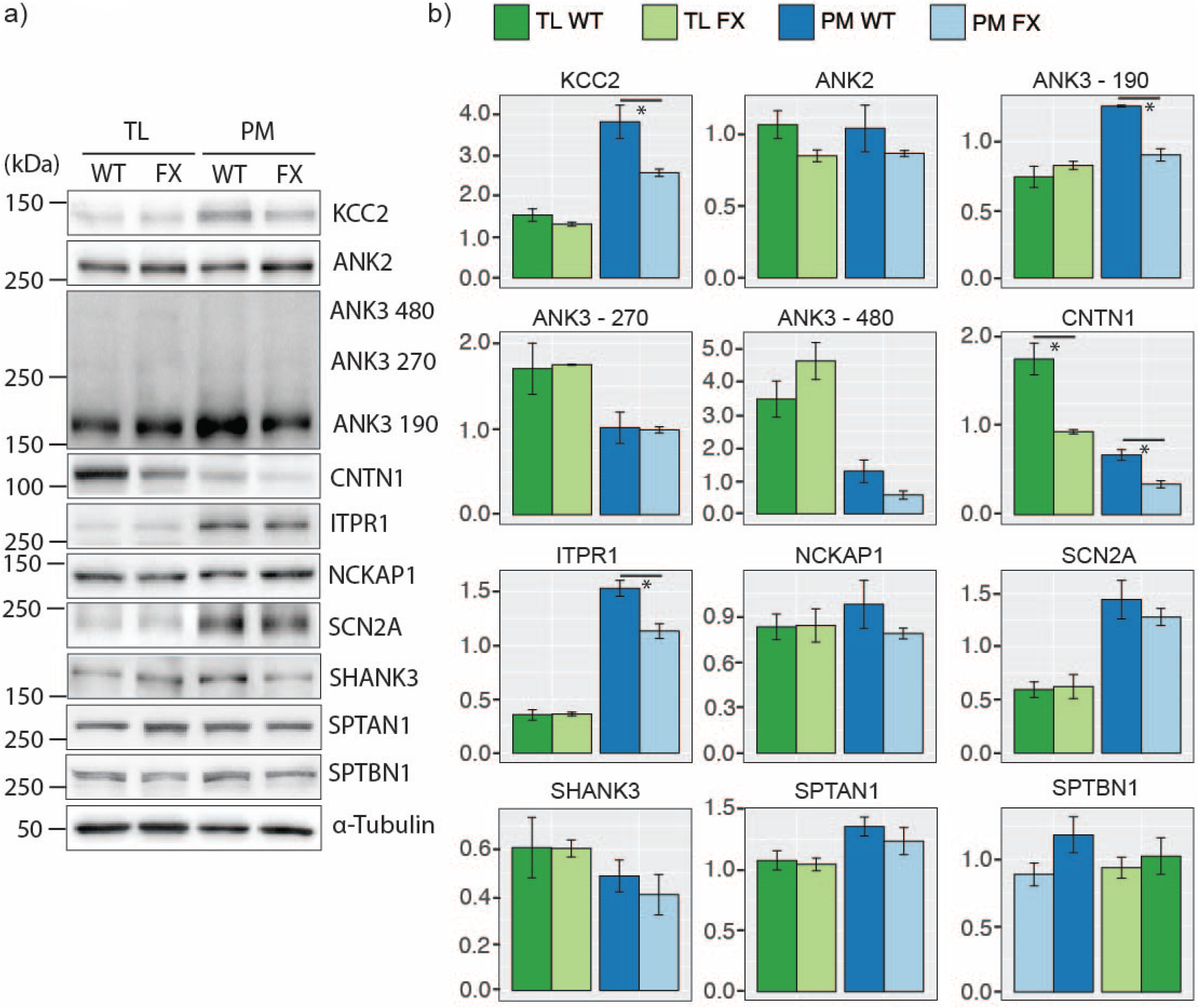
The total and plasma membrane expression levels of KCC2 and several associated ASD/Epi risk gene products are changed in FMR1 KO (FX) animals. **A.** Total forebrain lysates and plasma membrane lysates were resolved by SDS-PAGE and immunoblotted for selected high risk ASD/Epi risk gene product; ANK2, ANK3, CNTN1, ITPR1, NCKAP1, SCN2A, SHANK3, SPTAN1 and SPTBN1. **B.** The expression levels for each protein were quantified using densitometry and normalized to α-Tubulin loading controls (n=3). KCC2 (p=0.037), ANK3-190 (p=0.0014), CNTN1 (p=0.011), and ITPR1 (p=0.016) were significantly reduced in plasma membrane fractions.

### FMRP inactivation compromises KCC2 phosphorylation

KCC2 expression levels, membrane trafficking and activity have been shown to be subject to modulation via multiple phosphorylation sites within the C-terminal cytoplasmic domain. Our proteomic studies have revealed that KCC2 is intimately associated with multiple targets of FMRP, many of which are scaffold/signaling molecules. Thus, we sought to analyze the effects of FMRP inactivation, and subsequent reduction of KCC2-associated protein expression, on global KCC2 phosphorylation. To do so, we immunoprecipitated KCC2 from plasma membrane fractions prepared from WT and FX mice as outlined in Figure 1. To limit dephosphorylation purified material was resolved by SDS-PAGE and visualized using Coomassie (Figure 8a). Major bands of 125 kDa were seen with material purified on KCC2 antibody, but not control IgG. The respective bands were excised, digested with trypsin, and analyzed by LC-MS/MS. LC-MS/MS confirmed the 125 kDa bands were highly enriched in KCC2 with an average of 884 and 915 peptides detected (n=4), which equates to coverage of 53% and 43%, for KCC2 from WT and FX tissue respectively (Figure 8b).

**Figure 8.**
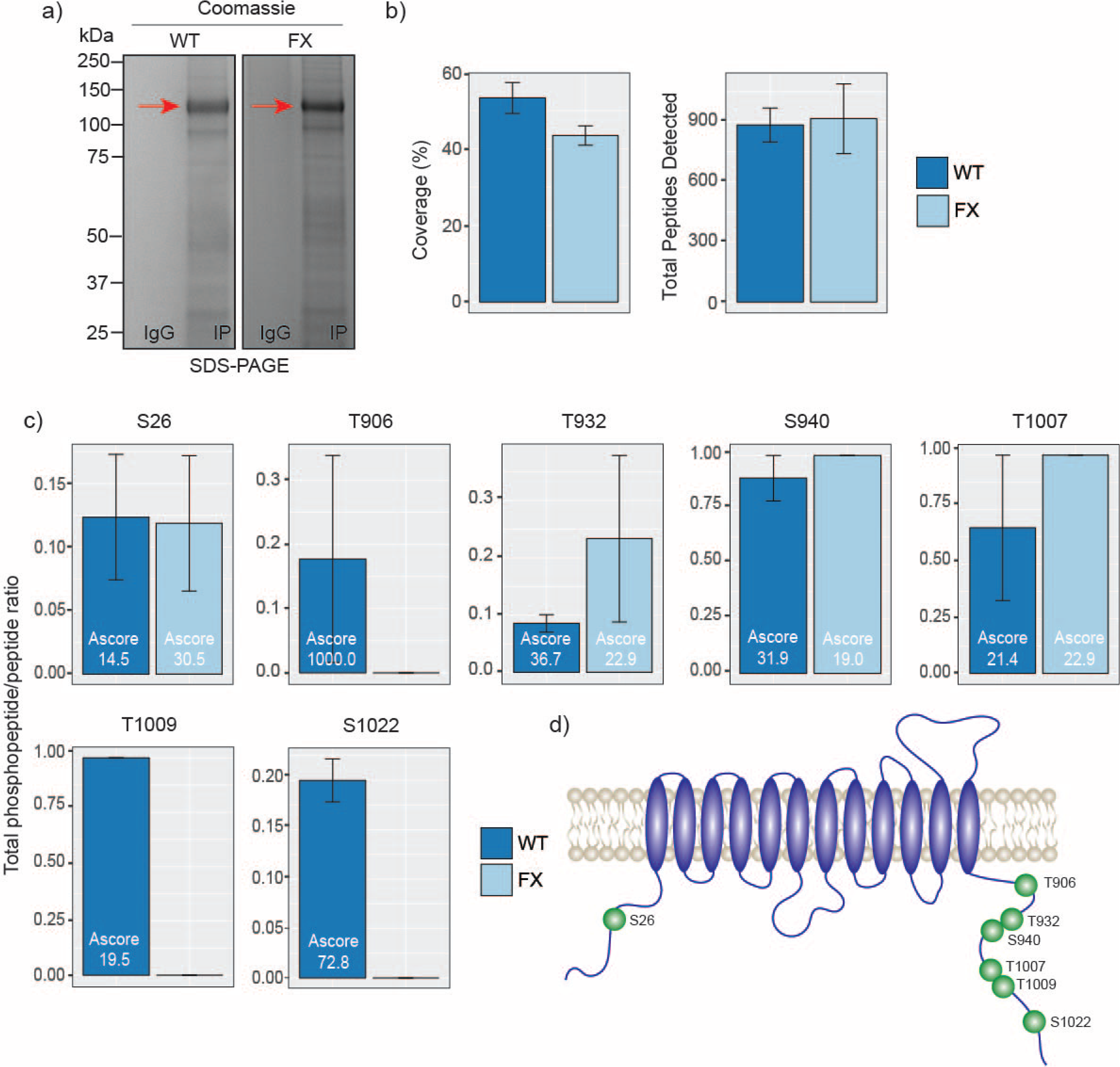
KCC2 is aberrantly phosphorylated in FMR1 KO (FX) mice. **A.** KCC2 was isolated from plasma membrane fractions of mouse forebrain and resolved by SDS-PAGE. **B.** Bar graphs of high confidence phosphorylated peptides relative to total detected peptides in wild type and FMR1 KO mice. For bars where phosphorylation was detected, A scores are superimposed to express phosphoryation confidence levels. **C.** Diagram of high confidence KCC2 phosphorylation site positions.

We identified 7 high confidence KCC2 phosphorylation sites based on A-scores greater than 19, which equates to 99% confidence in assignment (Figure 8c and d). One site (S26), is located on the N-terminus of KCC2, whereas the other 6 sites were located on the C-terminus (T906, T932, S940, T1007, T1009, S1022). We normalized the number of phosphorylated peptides detected for each site to the total number of peptides detected in that region and compared the ratios between KCC2 isolated from WT and FX. In this way we could compare changes in global KCC2 phosphorylation between WT and FX mice. This approach revealed that inactivation of FMRP resulted in the specific dephosphorylation of a subset of residues and generated insights into global phosphorylation changes in KCC2. Two of the well characterized sites, S940 and T1007, showed no changes in phosphorylation, along with two of the novel sites detected; S26, and T932. In contrast T906, T1009, and S1022, which are present in WT tissue, were completely absent in FX tissue (n=4). Collectively these results demonstrate that FMRP regulates the phosphorylation of a subset of residues within the C-terminal intracellular domain of KCC2, likely due to its control of the expression of several KCC2-interacting proteins.

## Discussion

### Isolation of stable multi-protein complexes enriched in KCC2

The K_+_/Cl_−_ co-transporter, KCC2, is expressed at the cell surface of mature neurons where it is of fundamental importance for maintaining intracellular chloride levels, and thereby the efficacy of neuronal inhibition. To gain insights into how neurons regulate the plasma membrane accumulation and activity of KCC2, we developed a method for the isolation of native multiprotein complexes from enriched in this transporter from brain plasma membranes. Following immuno-affinity purification, BN-PAGE and LC-MS/MS we were able to isolate highly purified native preparations of KCC2, which exhibited major molecular mass species between 300-800 kDa. This methodology provided very high sequence coverage for the intracellular and extracellular domains of KCC2, maximizing the probability detecting associated proteins and post-translational modifications.

KCC2 consists of 1139 amino acids including the signal sequence, which equates to a mass of 126 kDa without post-translational modifications. KCC2 is, however, subject to extensive post-translational modification including glycosylation at six sites so is often observed in a band of around 140 kDa (39). Previous work has shown extensive aggregation of KCC2 following SDS-PAGE that results in a band at 250 kDa, but this depends on whether it is heterologously expressed or the manner in which it is prepared from neuronal tissues (40). Indeed, we see no aggregation in our samples following SDS-PAGE, possibly due to the use of low detergent concentrations and samples immediately processed for SDS-PAGE without freeze thaw cycles. Interestingly, when resolved by BN-PAGE, KCC2 protein complexes form distinct bands at 300 kDa, 600 kDa and 800 kDa. The 300 kDa band contains approximately 6-fold more peptides for KCC2 than the next most abundant protein, indicating that this band consists largely of KCC2. At 300 kDa, this would indicate that in the plasma KCC2 exists as homodimers, which is consistent with recently published studies on recombinant KCC2 molecules visualized using Cryo-EM (39) Interestingly, the next most abundant protein in the 300 kDa species was KCC1, potentially indicating that KCC1/KCC2 heterodimers are present in the brain, albeit at lower levels than KCC2 homodimers. Consistent with this notion studies in oocytes suggest that KCC2 can form heterodimers with other KCCs (41). Whether these potential heterodimers are functional is unclear but are an exciting focus for future research.

The 600kDa and 800 kDa bands also contained KCC2, together with a range of other proteins. Significantly the ratio of these associated proteins relative to KCC2 increased with molecular mass. Given the stringency of our methodology and the use of native conditions these species represent stable-protein complexes that primarily reside in the plasma membrane. The proteins detected by a previous proteomic analysis of KCC2 complexes (42) overlapped with 13% and 16% of the proteins we detected in the 600 kDa and 800 kDa bands respectively. This low degree of overlap is likely due to the methodological differences between the studies; the previous work extracted KCC2 complexes from a crude membrane preparation and used on-bead digest prior to LC-MS/MS, rather than complexes purified from a plasma membrane fraction resolved by BN-PAGE, then distinct bands excised for analysis. The previously presented work is possibly more likely to detect transient interactions due to less sample processing. However, the method presented here ensured a sample of reduced complexity was analyzed by LC-MS/MS, resulting in a higher resolution snapshot of the proteins associated with high affinity with KCC2 on the neuronal plasma membrane (43, 44). Similarly to Mahadevan *et al*., we also did not detect NETO2 (45), GRIK2 (Gluk2) (46), EPB41L1 (4.1n) (47), ARHGEF7 (Beta-pix) (48), RCC1 (49) or with the signal transduction molecules; PKC, WNK, SPAK or OSR (8, 50). This could be either due the transient or low affinity nature of these interactions, or that these interactors are less abundant in complex with KCC2 than the interactors presented here, as unbiased methodologies provide information on associated proteins in rank order of abundance. LC-MS/MS further confirmed the veracity of our purification strategy as these high molecular weight species contained GABABRs, which has been previously shown to be associated with KCC2 (35). Interestingly we identified gephyrin associated with KCC2 in the 800 kDa band along with CYFIP1/2, NCKAP1 and CNTN1 and other proteins that are enriched at inhibitory synapses (51, 52). NLGN3 was also detected, which can be present at both inhibitory and excitatory synapses (53). KCC2 has previously been shown to be in close proximity to excitatory synapses and has been established to be associated with proteins enriched at these structures (42, 54, 55). In agreement with this we detected well characterized excitatory synapse proteins such as SHANK3, MYO5A, and DLG1 in our purifications (56–58).

### KCC2 is associated with a subset of highly interconnected ASD/Epi gene products

The most striking feature of the KCC2 proteome is that it is highly enriched for protein products of genes associated with ASD/Epi. Perturbation of KCC2 itself is associated with autism (59) and epilepsy (14, 15, 17, 59). Of the KCC2-associated proteins, ANK2, SHANK3, and SCN2A are all ranked in the highest group of ASDs by SFARI, mutations in the latter are also a cause of Dravet syndrome (38). We selected these proteins for further investigation, along with NCKAP1 (SFARI group 2), ANK3 (SFARI Group 3), ITPR1 (SFARI Group 4 and epilepsy risk gene), CNTN1 – an *in silico* predicted ASD (60, 61), SPTAN1 (highly implicated in epilepsy) (62), and SPTBN1 (indirectly genetically implicated in ASD) (63). These proteins were selected due to a combination of their disease implications and the number of detected peptides. They were also selected, as they are closely related to each other, including the well-established interactions between ankyrins and spectrins (64), which can recruit SCN2A at the cell surface (65). CNTN1 is also thought to interact with spectrins either directly or via CASPR1 (66). NCKAP1 is a part of the WAVE regulatory complex (WRC) that control actin dynamics (67). This complex can interact with ankyrins and SHANK3 (68).

### FMRP inactivation compromises KCC2 phosphorylation

Many of the components of the KCC2 proteome are targets of FMRP. Therefore, to assess the significance of this protein network on KCC2, we assessed the effects of FMRP inactivation on KCC2 phosphorylation, a key regulator of transporter activity. Plasma membrane levels of the FMRP targets proteins that are also KCC2 binding partners; ANK3, CNTN1 and ITPR1 were decreased in FX mice as were those of KCC2. In WT mice KCC2 was phosphorylated on up to seven residues including T906, S940, T1007. These play a critical role in regulating KCC2 function (13, 69). In addition to this we detected phosphorylation of S1022 consistent with a recent publication (39).

We also identified 3 further sites of phosphorylation; S26, T932 and T1009 respectively (70). Significantly, inactivation of FMRP abolished T906, T1009, and S1022 phosphorylation, without impacting on that of S26, T932, S940 or T1007. It is of note that these three phosphorylation sites all lie within the intracellular C-terminus of KCC2, and while T906 phosphorylation has been established to inhibit KCC2 activity, the roles that T1009 and T1022 phosphorylation play in modulating transporter function, protein-protein interactions, and/or membrane trafficking is unknown. Interestingly, T1009 and T1022 lie adjacent to, or within the isotonic domain of KCC2, suggesting that their phosphorylation may impact on its basal activity. Thus, it is tempting to speculate that modified KCC2 phosphorylation may contribute to aberrant GABAergic signaling that is central to the pathophysiology of FX and other ASDs.

In conclusion we have demonstrated that KCC2 is a point of convergence for multiple ASD/Epi risk gene products, which may act in part to coordinate its phosphorylation. Therefore, therapies targeting KCC2 may have broad utility in alleviating autism and associated epilepsies with varying genetic etiologies.

### Experimental Procedures

Unless otherwise stated chemicals were obtained from Sigma-Aldrich, St. Louis, MO, USA.

#### Animals

Animal studies were performed according to protocols approved by the Institutional Animal Care and Use Committee of Tufts Medical Center. 8-12-week-old male and female mice were kept on a 12-hour light/dark cycle with *ad libitum* access to food and water.

#### Antibodies

The following antibodies were used for immunoprecipitation (IP), immunoblot (IB), or immunocytochemistry (ICC): ANK2 (mouse, ICC, Invitrogen 33-3700), ANK2 (rabbit, IB, Bioss BS-6967R-TR), ANK3 (mouse, ICC, Neuromab 75-146), ANK3 (rabbit, IB, Synaptic Systems 386003), CNTN1 (rabbit, IB/ICC, Abcam Ab66265), ITPR1 (rabbit, IB/ICC, Alomone ACC-019), KCC2 (mouse, IP/ICC, Neuromab 75-013), KCC2 (rabbit, IB/ICC, Millipore 07-432), NCKAP1 (rabbit, IB, Abcam 126061), NCKAP1 (rabbit, ICC, Sigma HPA020449), SCN2A (mouse, ICC, Neuromab 75-024), SCN2A (rabbit, IB, Alomone ASC-002), SHANK3 (rabbit, IB/ICC, Alomone APZ-013), SPTAN1 (mouse, ICC, Abcam Ab11755), SPTAN1 (rabbit, IB, Cell Signaling 21225), SPTBN1 (rabbit, IB/ICC, Abcam Ab72239), α-Tubulin (mouse, IB, Sigma T9026).

Immunoblotting Sodium dodecyl sulphate polyacrylamide gel electrophoresis (SDS-PAGE) was carried out as previously described (71). Briefly, proteins were isolated in RIPA lysis buffer (50 mM Tris, 150 mM NaCl, 0.1% SDS, % sodium deoxycholate and 1% Triton X-100, pH 7.4) supplemented with mini cOmplete protease inhibitor and PhosSTOP phosphatase inhibitor tablets. Protein concentration was measured using a Bradford assay (Bio-Rad, Hercules, CA, USA). Samples were diluted in 2x sample buffer and 20 μg of protein was loaded onto a 7%, 10%, or 12% polyacrylamide gel depending on the molecular mass of the target protein. After separation by SDS-PAGE, proteins were transferred onto nitrocellulose membrane. Membranes were blocked in 5% milk in tris-buffered saline 0.1% Tween-20 (TBS-T) for 1 hour, washed with TBS-T, and then probed with primary antibodies diluted in TBS-T (dilution and incubation time dependent on the antibody). The membranes were washed and incubated for 1 hour at room temperature with HRP-conjugated secondary antibodies (1:5000 – Jackson ImmunoResearch Laboratories, West Grove, PA, USA). Protein bands were visualized with Pierce ECL (ThermoFisher) and imaged using a ChemiDoc MP (Bio-Rad). Band intensity was compared to α-tubulin as a loading control.

#### BN-PAGE

For blue native poly-acrylamide gel electrophoresis (BN-PAGE), proteins were isolated using Triton lysis buffer (150mM NaCl, 10mM Tris, 0.5% Triton X-100, pH 7.5). Samples were diluted in 4x NativePAGE sample buffer and G250 additive. Samples were loaded onto 4-16% NativePAGE gels and run using a mini gel tank (ThermoFisher, Waltham, MA, USA). Gels were run for approximately 2 hours and then prepared for coomassie staining or immunoblotting. For coomassie staining the gels were fixed in 50% ethanol and 10% acetic acid, washed in 30% ethanol, washed in water, then stained with EZ blue stain. The gels were destained in ultrapure water, imaged using a ChemiDoc MP (Bio-Rad), and bands excised for liquid chromatography tandem mass spectrometry (LC-MS/MS). For immunoblotting, proteins were transferred to PVDF membrane overnight. The membranes were then fixed in 8% acetic acid, washed with ultrapure water and air-dried before being briefly destained with 100% methanol. The membranes were then blocked, immunoblotted, and imaged as normal.

#### Plasma membrane isolation

Plasma membrane isolation from mouse forebrain was carried out using a modified version of a previously described method (72). Briefly, mouse forebrain was rapidly isolated and placed in ice-cold starting buffer (225mM mannitol, 75mM sucrose, 30mM Tris-HCl (pH 7.4). The tissue was then transferred to ice-cold isolation buffer (225mM mannitol, 75mM sucrose, 0.5% (wt/vol) BSA, 0.5mM EGTA, 30mM Tris-HCl, pH 7.4) supplemented with mini cOmplete protease inhibitor and PhosSTOP. 5-7 brains were used per immunoprecipitation. The brains were homogenized in the isolation buffer using 14 strokes of a Dounce homogenizer. The samples were transferred to centrifuge tubes for centrifugation, which was carried out at 4°C throughout. The samples were initially spun at 800xg for 5 minutes to remove nuclei and non lysed cells. The pellet was discarded, and the supernatant was spun again at 800xg for 5 minutes to remove residual nuclei and non lysed cells. The supernatant was transferred to high speed centrifuge tubes and spun at 10000xg for 10 minutes to remove mitochondria. The pellet was discarded, and the supernatant spun again at 10000xg for 10 minutes to remove mitochondrial contamination. The supernatant was then spun at 25000xg for 20 minutes to pellet plasma membranes. The pellet was resuspended in starting buffer and spun again at 25000xg for 20 minutes to remove cytosolic and ER/Golgi contamination. Finally, the plasma membrane fraction was purified on a discontinuous sucrose gradient (53%, 43%, 38% sucrose), and spun at 93000xg for 50 minutes to remove residual contamination from cytosol and other membranous compartments.

#### Protein purification

Protein G Dynabeads (ThermoFisher) were washed 3 times with phosphate buffered saline with 0.05% Tween-20 (PBS-Tween). The beads were resuspended in PBS-Tween and incubated overnight at 4°C with antibodies for the target protein at an experimentally predetermined bead:antibody ratio (Figure S1). The antibody was crosslinked onto the beads by washing twice with 0.2M triethanolamine (pH 8.2) (TEA), and then incubated for 30 minutes with 40mM dimethyl pimelimidate (DMP) in TEA at room temperature. The beads were transferred to 50mM Tris (pH7.5) and incubated at room temperature for 15 minutes. The beads were then washed 3 times with PBS-Tween and resuspended in solubilized plasma membranes in ice cold Triton lysis buffer, supplemented with mini cOmplete protease inhibitor and PhosSTOP. The immunoprecipitation reaction was incubated overnight at 4°C. The beads were then washed 3 times with PBS-Tween and eluted either with 2x sample buffer (for SDS-PAGE) or soft elution buffer (0.2% (w/V) SDS, 0.1% (V/V) Tween-20, 50 mM Tris-HCl, pH=8.0) (30) (for BN-PAGE).

#### Protein and phosphosite identification

Excised gel bands were cut into 1 mm_3_ pieces and subjected to modified in-gel trypsin digestion, as previously described (73). Briefly, gel pieces were washed and dehydrated with acetonitrile for 10 minutes and then completely dried in a speed-vac. Gel pieces were rehydrated with 50 mM ammonium bicarbonate solution containing 12.5 ng/μl modified sequencing-grade trypsin (Promega, Madison, WI, USA) and incubated for 45 minutes at 4°C. The excess trypsin solution was removed and replaced with 50 mM ammonium bicarbonate solution. Samples were then incubated at 37°C overnight. Peptides were extracted by washing with 50% acetonitrile and 1% formic acid. The extracts were then dried in a speed-vac (~1 hour). The samples were stored at 4°C until analysis. Before analysis the samples were reconstituted in 5 - 10 μl of HPLC solvent A (2.5% acetonitrile, 0.1% formic acid). A nano-scale reverse-phase HPLC capillary column was created by packing 2.6 μm C18 spherical silica beads into a fused silica capillary (100 μm inner diameter x ~30 cm length) with a flame-drawn tip (74). After equilibrating the column each sample was loaded via a Famos auto sampler (LC Packings, San Francisco, CA, USA) onto the column. A gradient was formed and peptides were eluted with increasing concentrations of solvent B (97.5% acetonitrile, 0.1% formic acid). As each peptide was eluted, they were subjected to electrospray ionization and then entered into an LTQ Orbitrap Velos Pro ion-trap mass spectrometer (Thermo Fisher Scientific, San Jose, CA, USA). Eluting peptides were detected, isolated, and fragmented to produce a tandem mass spectrum of specific fragment ions for each peptide. Peptide sequences (and hence protein identity) were determined by matching protein or translated nucleotide databases with the acquired fragmentation pattern using Sequest (ThermoFinnigan, San Jose, CA, USA) (75). For phosphopetide detection the modification of 79.9663 mass units to serine, threonine, and tyrosine was included in the database searches. Phosphorylation assignments were determined by the A score algorithm (76). All databases include a reversed version of all the sequences and the data was filtered to between a one and two percent peptide false discovery rate.

#### Primary neuron culture

Mouse cortical and hippocampal mixed cultures were created from P0 mouse pups as previously described (77). Briefly, P0 mice were anesthetized on ice and the brains removed. The brains were dissected in Hank’s buffered salt solution (Invitrogen/ThermoFisher) with 10 mM HEPES. The cortices and hippocampi were trypsinized and triturated to dissociate the neurons. Cells were counted using a hemocytometer and plated on poly-l-lysine-coated 13mm coverslips in 24-well plate wells at a density of 2 × 10_5_ cells/ml in Neurobasal media (Invitrogen/ThermoFisher). At days *in vitro* (DIV) 18, cells were fixed in 4% paraformaldehyde in PBS for 10 minutes at room temperature. They were then placed in PBS at 4°C until being processed for immunocytochemistry.

#### Immunocytochemistry

Fixed primary neurons were permeabilized for 1 hour in blocking solution (3% BSA, 10% normal goat serum, 0.2 M glycine in PBS with 0.1% Triton-X100). Cells were exposed to primary and then fluorophore-conjugated secondary antibodies diluted in blocking solution for 1h each at room temperature. The coverslips were then washed in PBS, dried, and mounted onto microscope slides with Fluoromount-G (SouthernBiotech, Birmingham, AL, USA). The samples were imaged using a Nikon Eclipse Ti (Nikon Instruments, Melville, NY, USA) or Leica Falcon (Leica Microsystems, Buffalo Grove, IL, USA) confocal microscope using a 60x oil immersion objective lens. Image settings were manually assigned for each fluorescent channel. For image processing, the background was subtracted for each fluorescent channel and the median filter was applied (Radius = 1 pixel) on Fiji Software (78). The line scans (white lines) used for protein localization were generated using the PlotProfile function in FIJI and represent the fluorescent intensity of single pixels against the distance of a manually drawn line (approximately 1 μm) on dendrites.

#### Densitometry

For Western blot analysis, bands from raw images were analyzed using the FIJI densitometry features. Biological replicates were run on the same gels for comparison, and area under the curve was calculated for each band. Average signal and standard error were calculated for each treatment group and ANOVA carried out using R for statistical comparison of protein expression levels.

#### Bioinformatics and Statistics

Detected peptides for each Uniprot ID were compared to the Uniprot mouse reference sequence for annotation using R. KCC2 peptide sequences were aligned to the mouse KCC2 reference sequence (Uniprot ID: Q91V14) using the Multiple Sequence Alignment (MSA) package in R (accessed January 10_th_, 2019). Proteins with significant affinity for IgG alone were removed according to (31, 32) along with proteins with fewer than 3 total peptides across 3 of 4 replicates. Total peptide counts for detected proteins in each protein band were compared across 4 biological repeats, Venn diagrams produced using the Vennerable package in R (accessed January 10_th_, 2019), and only proteins detected in all repeats considered for downstream analysis. These total peptide counts for the lists of proteins contained within each gel band were normalized by z-transformation and used for Principle Component Analysis (PCA), which was carried out using the PCA functions in R. The protein lists for each band were ordered according to total peptide counts, a measure for relative abundance, and the top 150 proteins used for network analysis. The protein lists were compared against the latest version of the StringDB database (34) to establish known interactions and the Gene Ontology (GO) terms for each protein (Table S3 and S4). The interaction for each protein with KCC2 was imputed using only high confidence, experimental evidence and network diagrams were constructed in R using the igraph package (accessed February 1_st_, 2019). A master list of epilepsy and autism risk genes was constructed from the SFARI database (29), the list of genes that are the target of FMRP (36), and the EpilepsyGene (37) database (Table S5). This data was overlaid onto the KCC2 proteome to assess the abundance of autism/epilepsy risk genes associated with KCC2.

## Acknowledgements

S.J.M. is supported by National Institutes of Health (NIH)–National Institute of Neurological Disorders and Stroke Grants NS051195, NS056359, NS081735, R21NS080064 and NS087662; NIH–National Institute of Mental Health Grant MH097446

## Conflict of interest statement

S.J.M serves as a consultant for AstraZeneca, and SAGE Therapeutics, relationships that are regulated by Tufts University. S.J.M holds stock in SAGE Therapeutics.

## Author Contributions

JLS and SJM conceptualized the project, analyzed the data and wrote the paper. GK performed immunocytochemistry experiments. CC and QR performed western blot experiments. MARS and CEB maintained the mouse colony and performed genotyping. TZD and NJB edited the paper.

